# Genetic architecture and functional consequences of lateral root length in maize (*Zea mays* L.)

**DOI:** 10.1101/2025.09.02.673713

**Authors:** Fabio Guffanti, Daniela Scheuermann, Claude Urbany, Stefan Reuscher, Thomas Presterl, Milena Ouzunova, Silvio Salvi, Caroline Callot, Chris-Carolin Schön

## Abstract

Understanding the genetic basis of root architecture and its relevance for crop productivity can contribute to the sustainable intensification of agriculture. Leveraging the phenotypic and allelic diversity of an Austrian maize landrace, we dissected the genetic basis of lateral root (LR) length across developmental stages. LR length, a relevant trait for breeding resource-efficient varieties, showed high heritability in our experiments. We discovered eight quantitative trait loci (QTL) for LR length at the reproductive stage R2, overlapping with four QTL at stage R6 but not with QTL detected at vegetative stage V6, suggesting that the genetic regulation of LR length might differ in vegetative and reproductive stages. We fine-mapped *qlr1*, the most significant QTL for LR length, to a region of 2.3 Mb containing 46 annotated genes. Based on whole-genome sequence and comparative genomics analyses we suggest a candidate gene underlying *qlr1*. Additionally, we examined the impact of nitrogen, phosphorus, and irrigation treatments on root and shoot development, finding that LR length positively correlates with biomass accumulation under optimal nutrient supply but not under nitrogen stress. Our work provides insights into the genetic regulation of LR length in maize and its relevance for the adaptation to specific growing environments.

**Highlight:** Multiple QTL affect lateral root length in maize at different developmental stages. The correlation between lateral root length and biomass accumulation varies under diverse field conditions.

## Introduction

Roots are critical to crop productivity and sustainability, as water and nutrient availability impair yield potential of all major crops, including maize (Mueller et al., 2012; Lynch, 2022). Breeding for root traits that enhance resource acquisition, particularly in low-fertility soils, is suggested to play an important role for the sustainable intensification of agriculture (Lynch, 2007). The maize root system comprises embryonic roots, namely the primary and seminal roots, and post-embryonic roots originating from the shoot (Hochholdinger et al., 2018). Shoot-borne roots (also called nodal roots) develop in consecutive shoot whorls (or nodes) and dominate the adult maize root system. They are classified as crown and brace roots depending on their development below or above the soil surface (Hochholdinger et al., 2004).

Lateral roots (LR), which emerge post-embryonically from all root types, constitute most of the total root length and exhibit high responsiveness to environmental stimuli, such as water, nutrients, and microbial interactions (Yu et al., 2014; Yu et al., 2016; Viana et al., 2022). In young maize plants, shoot-borne roots and their LRs already constitute most of the total root length and take up the majority of the water (Singh et al., 2010; Ahmed et al., 2018). They reach their maximum cumulative length between flowering time and two weeks after flowering, corresponding to the developmental stage R2 in field-grown maize (Peng et al., 2010; Peng et al., 2012), providing anchorage and facilitating soil resource acquisition throughout the plant’s life cycle (Hochholdinger et al., 2004).

LR length, branching frequency, and plasticity are important breeding targets to improve productivity, especially under sub-optimal growing conditions. Maize recombinant inbred lines differing in LR length and branching density at flowering have been shown to exhibit differential biomass accumulation and yield in the field under irrigation, nitrogen, and phosphorus treatments. Genotypes with few and long LRs performed better under low water and nitrogen availability, while those with short and dense LRs performed better in low phosphorus conditions (Zhan and Lynch, 2015; Zhan et al., 2015; Jia et al., 2018).

In maize, genes affecting LR initiation, elongation, and number have been described for primary and seminal roots (Woll et al., 2005; von Behrens et al., 2011; Lu et al., 2019; Gautam et al., 2021; Baer et al., 2023; Zhang et al., 2023). In addition, the number of QTL mapping studies for shoot-borne root traits has increased in recent years, identifying QTL of small to moderate effect, and suggesting a quantitative genetic architecture. These studies have primarily focused on axial roots (Zhang et al., 2018c; Wang et al., 2021; Li et al., 2024), while LR traits such as length and density have been explored less (Schneider et al., 2020). A valuable resource to study maize root traits is the genetic diversity offered by landraces, which might harbor new, favourable alleles compared to elite material (Mayer et al., 2020). A library of doubled-haploid (DH) lines derived from three European maize landraces showed striking genetic diversity for many agronomic traits relevant for breeding (Mayer et al., 2017; Hoelker et al., 2019). This natural mutant collection was successfully used to identify a QTL related to early plant development, including the discovery of the underlying causal gene and elucidation of its functional consequences and allelic diversity (Urzinger et al., 2025). In the present work, we investigated the natural variation of shoot-borne root architecture in DH lines derived from the Austrian maize landrace “Kemater Landmais Gelb”. We developed a series of extensively characterized bi-parental populations derived from two DH lines with large differences in their LR length and similar genomic background. The objectives of this study were to uncover the genetic architecture of LR length in shoot-borne roots during vegetative and reproductive stages, fine-map and characterize the most significant QTL for LR length, and evaluate the impact of LR length on shoot biomass accumulation under different phosphorus, nitrogen and water regimes.

## Material and Methods

### Plant Material

In previous work, a library of more than 1000 doubled-haploid (DH) lines was generated sampling a large number of individuals from three European maize landraces (Mayer et al., 2022). They were chosen to represent the molecular diversity of a comprehensive panel of 35 flint landraces, covering a broad geographical region of Europe (Mayer et al., 2017). A detailed description of the development, molecular and phenotypic characterization of the landrace-derived DH library is given in Hoelker et al. (2019). Briefly, DH lines were genotyped with the 600k Affymetrix® Axiom® Maize Genotyping Array (Unterseer et al., 2014), hereinafter referred to as the 600k array and phenotyped for agronomic traits in up to 11 field environments. To generate bi-parental QTL mapping populations for adult root traits, we analysed the mature root system (developmental stage R2) of a subset of 18 DH lines from one of the three landraces (Kemater Landmais Gelb, KE) in preliminary field trials in Freising (FS, Germany) and Bologna (BOL, Italy) in 2020. The 18 DH lines formed 9 pairs, each sharing a similar genomic background but carrying contrasting haplotypes associated with lodging and/or seedling root traits in earlier genome-wide association studies (GWAS) (Mayer et al., 2020; Guffanti et al., unpublished). One pair of these DH lines, KE0413 (P1) and KE0113 (P2), with similar genomic background (74,978 polymorphic SNPs out of 501,124 SNPs), similar flowering time, and contrasting lateral root (LR) length, was crossed to generate the mapping populations used in this study, comprising F_2_ individuals, recombinant inbred lines (RILs) and heterogeneous inbred families (HIFs) (**Figure 1**).

**Figure 1.**
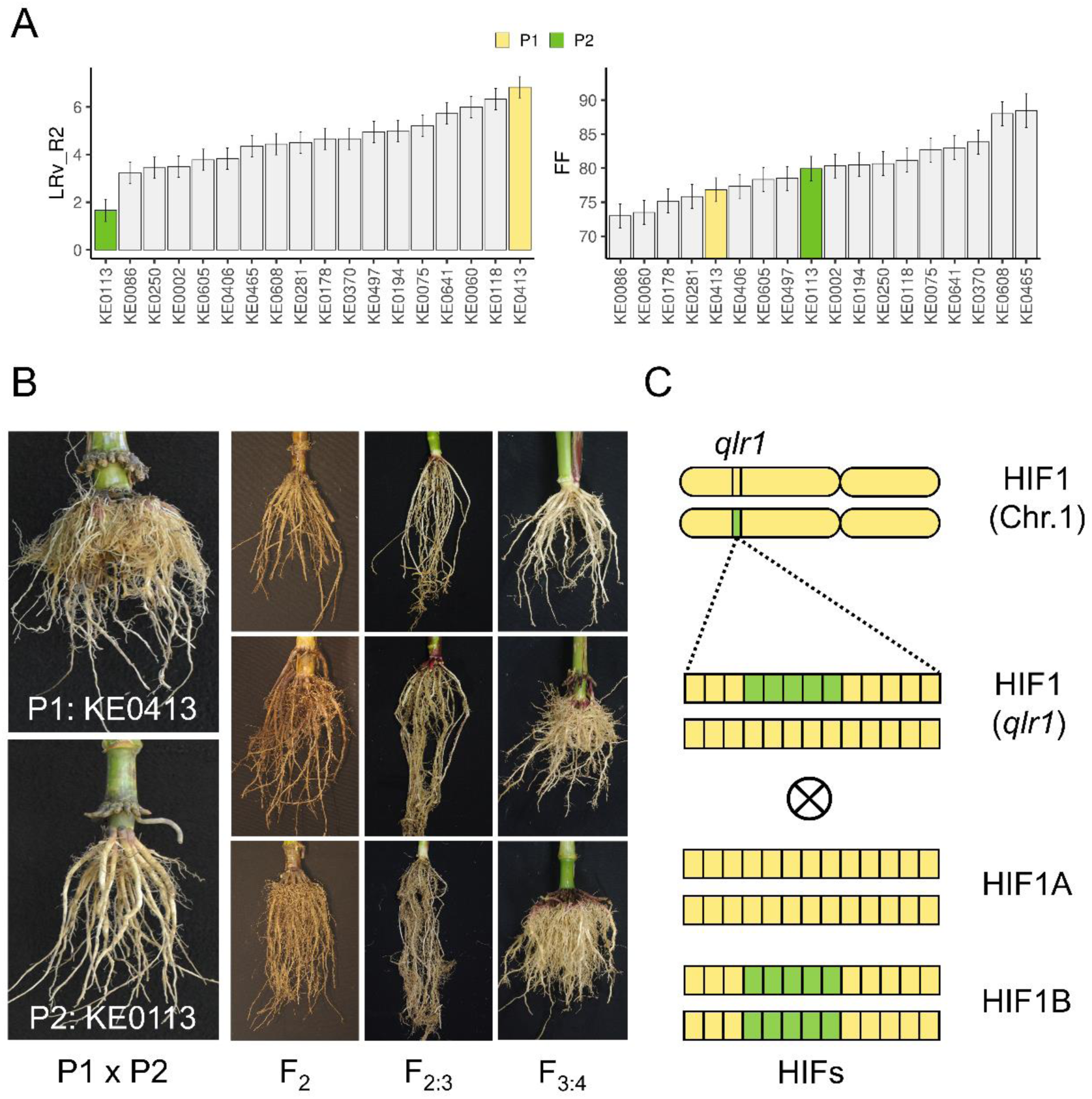
Selection of parental DH lines and development of plant material. **A)** LR length at stage R2 (LRv_R2) and flowering time (FF) of 18 landrace-derived DH lines evaluated in experiment E0. Bars represent adjusted means across two locations ± standard errors. **B)** P1 (KE0413) and P2 (KE0113) were crossed to develop QTL mapping populations segregating for LR length assessed on F_2_ individuals (stage R6, E1), F_2:3_ (stage V6, E2) and F_3:4_ progenies (stage R2, E3). **C)** Development of one HIF (HIF1) used for fine-mapping *qlr1.* HIFs were derived by self-pollination of F_3_, F_4_, or F_5_ individuals with heterozygous fragments of *qlr1*. Each resulting HIF is represented by two contrasting lines (HIFA, HIFB) homozygous for the P1 or the P2 allele (yellow = P1 allele, green = P2 allele). Vertical segments indicate schematically the position of the 14 KASP markers used to genotype the HIFs. HIFs differ with respect to the size and position of contrasting fragments.

### Development of recombinant inbred lines and heterogeneous inbred families

The F_2_ segregating population comprised 554 individuals. Through self-pollination, 150 F_2:3_ and 285 F_3:4_ RILs were derived from F_2_ and F_3_ individuals, respectively. Genotyping was performed on F_2_ and F_3_ individuals using a custom 15K SNP Illumina array proprietary to KWS SAAT SE & Co. KGaA (hereinafter referred to as 15K SNP array).

For QTL mapping F_2_ individuals as well as F_2:3_ and F_3:4_ RILs were used.

Based on the results of the QTL mapping experiments (**Figure 2**), the most significant QTL for LR length on Chr. 1 (*qlr1*) was targeted for fine-mapping. 14 KASP markers polymorphic between the parental lines and spanning a 26 Mb interval on Chr.1 (from AX-91333740, pos. 30671047 on the B73v5 physical map, to AX-90529307, pos. 56348234) were synthesised based on the 600K array probe sequences to identify recombination breakpoints within *qlr1*. Individual F_3_, F_4_, or F_5_ plants carrying different heterozygous fragments of *qlr1* at the genotyped KASP markers were self-pollinated to generate 43 HIFs (**Figure 1C**). The resulting progenies, segregating for the marker alleles of the respective fragments in a near-isogenic background, were examined in the fine-mapping experiments, either directly or following seed multiplication.

**Figure 2.**
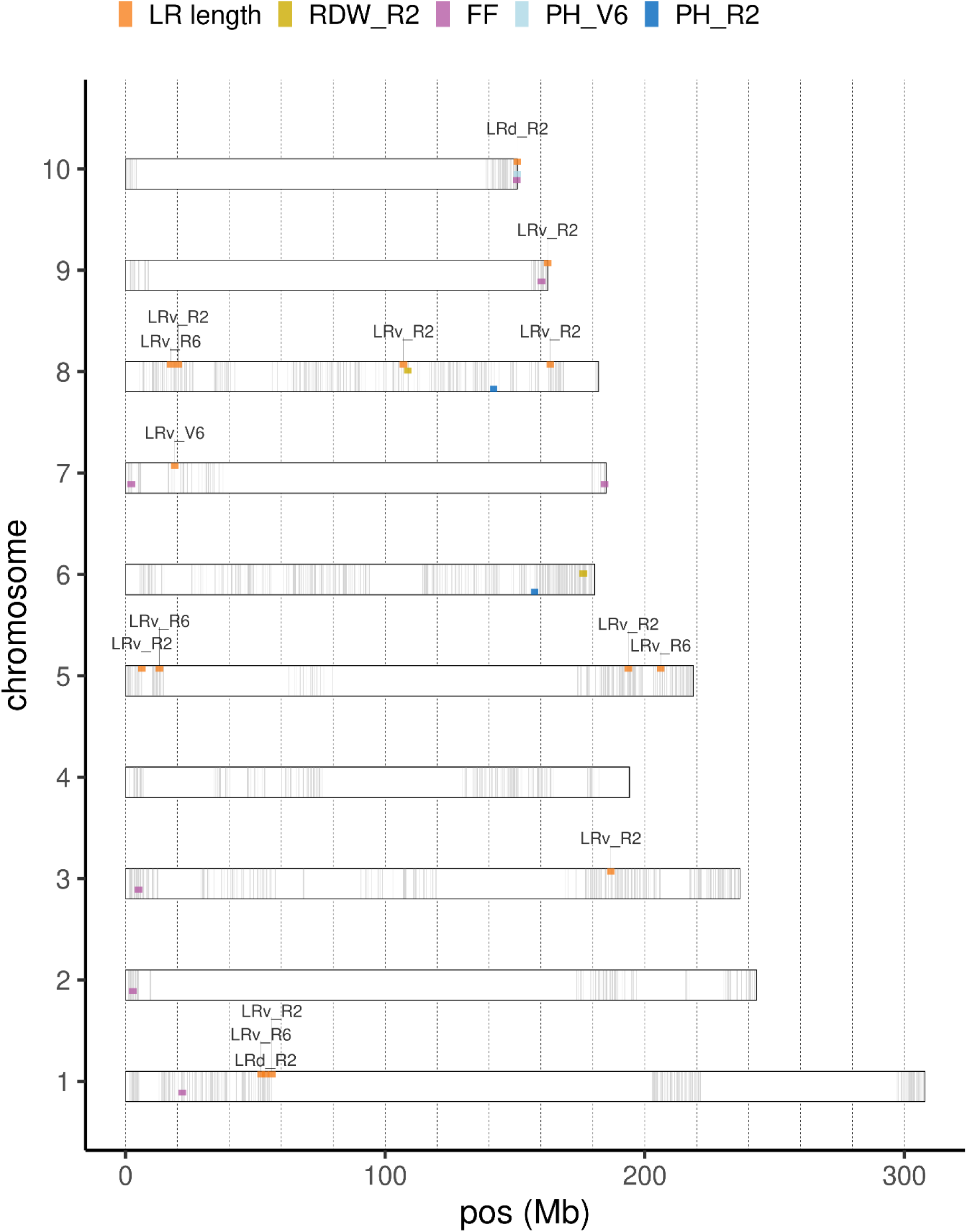
QTL mapping for root and agronomic traits. QTL positions on the B73 v5 physical map for traits lateral root (LR) length assessed by visual scoring at developmental stages V6, R2, and R6 (LRv_V6, LRv_R2, LRv_R6) and using DIRT at stage R2 (LRd_R2), root dry weight at stage R2 (RDW_R2), flowering time (FF), and plant height at stages V6 and R2 (PH_V6, PH_R2). Rectangles represent the 10 chromosomes, grey segments indicate polymorphic markers between P1 and P2 used for genetic map construction and QTL mapping. Coloured points denote QTL associated with different traits. QTL for LR length are labelled.

Two F_4_ individuals homozygous for the P1 allele at all 14 KASP markers within *qlr1* and two F_4_ individuals homozygous for the P2 allele for 13 KASP markers within *qlr1* (from 34 Mb to 56 Mb) were self-pollinated twice to generate F_4:6_ RILs contrasting for the entire region of *qlr1* to be included in the fine-mapping experiments.

### Growth conditions and experimental design

A total of eight experiments (E0-E7) are presented in this study. In E0, the aim was to screen the adult root architecture of DH lines from the landrace KE to select the parents of the bi-parental populations. In E1-E3 the aim was to map QTL for LR length at different developmental stages. In E4-E7, the aim was to fine-map and characterize the effect of *qlr1*, the most significant of the QTL identified for LR length.

Details for all experiments with respect to plant material, treatments, and replications are summarized in **Table 1**.

**Table 1.**
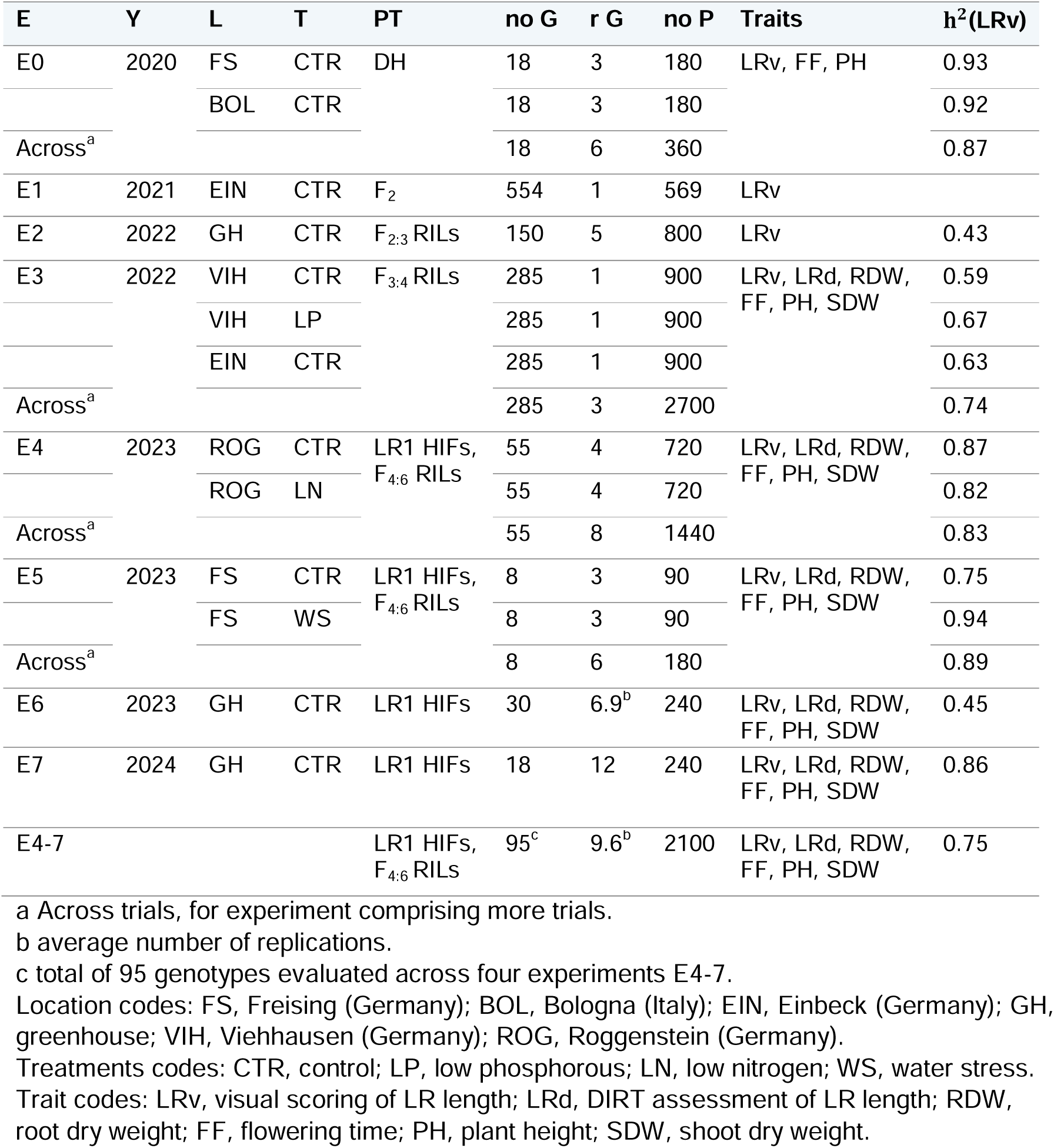
Detailed information on experiments (E) E0-E7. Given are the year (Y) and location (L) of evaluation, applied treatment (T), progeny type (PT), number of genotypes (no G), number of replications of the genotypes (r G), total number of plants measured including genotypes and checks (no P), Traits with significant genetic variance and entry mean heritability of visually scored LR length [h^2^ (LRv)].

| E | Y | L | T | PT | no G | r G | no P | Traits | $h^2$(LRv) |
| --- | --- | --- | --- | --- | --- | --- | --- | --- | --- |
| E0 | 2020 | FS | CTR | DH | 18 | 3 | 180 | LRv, FF, PH | 0.93 |
|  |  | BOL | CTR |  | 18 | 3 | 180 |  | 0.92 |
| Across<sup>a</sup> |  |  |  |  | 18 | 6 | 360 |  | 0.87 |
| E1 | 2021 | EIN | CTR | F<sub>2</sub> | 554 | 1 | 569 | LRv |  |
| E2 | 2022 | GH | CTR | F<sub>2:3</sub> RILs | 150 | 5 | 800 | LRv | 0.43 |
| E3 | 2022 | VIH | CTR | F<sub>3:4</sub> RILs | 285 | 1 | 900 | LRv, LRd, RDW, FF, PH, SDW | 0.59 |
|  |  | VIH | LP |  | 285 | 1 | 900 |  | 0.67 |
|  |  | EIN | CTR |  | 285 | 1 | 900 |  | 0.63 |
| Across<sup>a</sup> |  |  |  |  | 285 | 3 | 2700 |  | 0.74 |
| E4 | 2023 | ROG | CTR | LR1 HIFs, F<sub>4:6</sub> RILs | 55 | 4 | 720 | LRv, LRd, RDW, FF, PH, SDW | 0.87 |
|  |  | ROG | LN |  | 55 | 4 | 720 |  | 0.82 |
| Across<sup>a</sup> |  |  |  |  | 55 | 8 | 1440 |  | 0.83 |
| E5 | 2023 | FS | CTR | LR1 HIFs, F<sub>4:6</sub> RILs | 8 | 3 | 90 | LRv, LRd, RDW, FF, PH, SDW | 0.75 |
|  |  | FS | WS |  | 8 | 3 | 90 |  | 0.94 |
| Across<sup>a</sup> |  |  |  |  | 8 | 6 | 180 |  | 0.89 |
| E6 | 2023 | GH | CTR | LR1 HIFs | 30 | 6.9<sup>b</sup> | 240 | LRv, LRd, RDW, FF, PH, SDW | 0.45 |
| E7 | 2024 | GH | CTR | LR1 HIFs | 18 | 12 | 240 | LRv, LRd, RDW, FF, PH, SDW | 0.86 |
| E4-7 |  |  |  | LR1 HIFs, F<sub>4:6</sub> RILs | 95<sup>c</sup> | 9.6<sup>b</sup> | 2100 | LRv, LRd, RDW, FF, PH, SDW | 0.75 |

a Across trials, for experiment comprising more trials.
b average number of replications.
c total of 95 genotypes evaluated across four experiments E4-7.
Location codes: FS, Freising (Germany); BOL, Bologna (Italy); EIN, Einbeck (Germany); GH, greenhouse; VIH, Viehhausen (Germany); ROG, Roggenstein (Germany).
Treatments codes: CTR, control; LP, low phosphorous; LN, low nitrogen; WS, water stress.
Trait codes: LRv, visual scoring of LR length; LRd, DIRT assessment of LR length; RDW, root dry weight; FF, flowering time; PH, plant height; SDW, shoot dry weight.

In E0, 18 DH KE lines and two flint inbred lines were evaluated in field trials in two locations (Bologna, Italy and Freising, Germany) in 2020 for root traits at stage R2 and agronomic traits throughout development. P1 and P2 were included as check genotypes in all subsequent experiments, together with three flint inbred lines in E1, E3, and E4. In E1, 554 F_2_ individual plants were evaluated in one field location (Einbeck, Germany) in 2021 for root traits at stage R6. In E2, 150 F_2:3_ progenies were evaluated in the greenhouse in 2022 for root and shoot traits at the vegetative stage V6. In E3, 285 F_3:4_ RILs were evaluated in three field trials in two locations (Viehhausen, Germany and Einbeck, Germany) in 2022 for root traits at stage R2 and agronomic traits throughout development. In Viehhausen, two phosphorus (P) treatments were applied in the same field with naturally low P content: in the control trial 100 kg P/ha were supplied, in the low P trial no P was supplied. In E4, a total of 55 progenies from the bi-parental cross comprising 25 HIFs (12 F_3:5_ and 13 F_4:6_), and 4 F_4:6_ RILs (contrasting for *qlr1*, two RILs with P1 alleles and two RILs with P2 alleles) were evaluated in two trials in the same field (Roggenstein, Germany) in 2023 for root traits at stage R2 and agronomic traits throughout development under two nitrogen (N) treatments. In the control trial, 170 kg N/ha were supplied, in the low N trial, no N was supplied. In E5, a subset of 8 progenies from the bi-parental cross tested in E4, consisting of 3 F_4:6_ HIFs, and 2 F_4:6_ RILs (contrasting for *qlr1*, one RIL with P1 alleles and one RIL with P2 alleles) were evaluated under two irrigation treatments in two trials in the same location (Freising, Germany) for root traits at stage R2 and agronomic traits throughout development. The water stress (WS) trial was performed in a rainout shelter, where the water supply was restricted to 138 l m^−2^ from May to September. The control trial was performed in a different field, exposed to rain, and irrigated when necessary. In E6, 13 F_3:4_ HIFs were evaluated in the greenhouse in 2023 for root traits at stage R2 and agronomic traits throughout development. In E7, 9 HIFs (7 F_4:6_ and 2 F_5:7_) were evaluated in the greenhouse in 2024 for root traits at stage R2 and agronomic traits throughout development. In the field experiments, plant protection, irrigation, herbicide, and fertilizer application were conducted according to standard agronomic practice for the respective soil types and locations unless specified otherwise. The experimental units were single or double-row (only E4) plots of 20 plants per 3 m row. In the greenhouse, the conditions were maintained at 25°C-35°C/18°C-20°C day/night, 40% relative humidity (RH), and a light intensity of 600 μmol m^-2^s^-1^ was supplied by ceramic metal-halide lamps in addition to the natural light. Experimental units were single plants grown in pots of 3 l (top diameter: 16 cm, height: 20 cm, up to stage V6) or of 15 l (top diameter: 33 cm, height: 22 cm, up to stage R2). The growing substrate consisted of a mixture of C-700 Stender (https://stender.de/) potting substrate (sieved with 10 mm mesh) and sand (0.7-1.2 mm) in a ratio of approximately 75/25 (v/v).

### Phenotypic data collection

We measured root and shoot traits at different developmental stages. For the assessment of root traits, the stem was severed at 10 cm from the surface level and the root stocks (three root stocks per plot for the field trials) were harvested and washed as described in Trachsel et al. (2011) and photographed with one or two excised roots from the last developed whorl with a black background. The root pictures were analysed with the software DIRT (Das et al., 2015), which computed 34 monocot root traits from the root stock and 11 traits from the excised roots. In addition, LR length was visually scored from the images of the root stock (LRv, 1-9 score, the scoreboard used for LRv is shown in **Figure S1**). Root stock fresh and dry weight (RFW, RDW g/plant) and water content (RWC, %) were evaluated on the same plants. In the fine-mapping experiments (E4-E7) root stocks were dissected. The number of seminal roots (N.SR, count) and the number of whorls where shoot-borne roots originated was counted (N.whorls, count). The number of shoot-borne roots reaching the ground was counted for all developed whorls separately and aggregated (N.SBR, count). From whorl 2 to whorl 7, three randomly selected excised roots per whorl were photographed with a black background, analysed with the software DIRT for the excised root traits, and visually scored for LR length.

The collected shoot traits included stand count (STC, count), early vigour score (EV, 1-9 score), plant height (PH, cm), leaf greenness measured with a chlorophyll-content-meter (SPAD-502Plus, Konica Minolta, Inc., Japan), (SPAD), male and female flowering (MF / FF, days after sowing), anthesis-silking interval (ASI, days), shoot fresh and dry weight (SFW, SDW, g/plant), shoot water content (SWC, %), grain fresh and dry yield (GFY, GDY, dt/ha), grain dry matter content (GDC, %).

If not stated otherwise, measurements in the field experiments were taken on three randomly chosen plants per plot. Scores refer to the entire plot. In the greenhouse, measurements were taken on single plants.

For detailed information on trait abbreviations and assessment in different experiments, refer to **Table S1**.

### Phenotypic data analysis

All statistical analyses were done in R (R Core Team, 2025). The statistical model used for estimating genotype and genotype by trial variance components was:

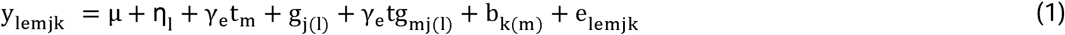

Where y_lemjk_ are the phenotypic observations, µ is the overall mean, η_l_ is the effect of the group (l=1,2 denotes checks and genotypes from the bi-parental cross, respectively), γ_e_ is a dummy variable with γ_e_ = 1 in analyses comprising experiments with several trials (E0, E3, E4, E5) and γ_e_= 0 otherwise, t_m_ is the effect of the trial m, g_j(l)_ is the effect of genotype j nested in group l, tg_mj(l)_ is the interaction of trial m and genotype j nested in group l, b_k(m)_ is the effect of block k nested in trial m, e_lemjk_ denotes the residual error. Phenotypic observations from plots with less than three plants were set to missing values. Outliers were manually curated by inspection of the residual plots.

To estimate variance components, g_j(l)_ was treated as fixed for l = 1, all other effects except η were treated as random and were assumed to be independent and identically normally distributed with mean zero and the respective variance component. Variance components and their standard errors were estimated using the restricted maximum-likelihood method implemented in ASReml-R package version 4 (Butler, 2017).

Trait heritabilities (h^2^) were calculated on an entry-mean basis according to Holland et al. (2003). Variance component estimates were considered significant when exceeding twice their standard errors.

Adjusted entry means within and across trials were obtained from equation (1), treating g_j(l)_ and t_m_ as fixed effects and dropping the term η_l_ from the model. The significance of the fixed effects was tested with an incremental Wald test implemented in ASReml-R. Fixed terms were considered significant for P < 0.05.

Phenotypic correlations among traits were calculated as the Pearson correlation coefficient of the adjusted means within and across trials for pairwise trait combinations. To account for multiple testing, we applied the Bonferroni–Holm correction (Holm, 1979). Correlations were considered significant for P < 0.05.

### Genotypic data analysis

The two parental lines, 550 F_2_ individuals phenotyped in E1, and 284 F_3_ founder individuals were genotyped using the 15K SNP array. SNPs not represented in the 600 K array, with missing calls in P1 or P2, heterozygous in P1 or P2, with > 5% missing calls, or with > 5% non-parental alleles were discarded. Of the SNPs overlapping with the 600K array, 97.4% and 96.7% were retained for the F_2_ and F_3_ datasets, respectively. Among the retained SNPs, 27.9% were polymorphic in the F_2_ and 17.6% in the F_3_, with 2212 overlapping between the two datasets. These overlapping SNPs were used for constructing genetic maps and for QTL mapping.

### Genetic maps and QTL mapping

Two genetic maps were generated based on the Haldane mapping function with genotypic data from F_2_ and F_3_ individuals using the package R/qtl (Arends et al., 2010). The marker order in both maps corresponded to their physical order on the B73 AGPv5 (Hufford et al., 2021) physical map (**Figure S2**). One F_3_ individual with an excessive number of crossing overs was excluded from QTL mapping.

For the F_2_ QTL mapping, phenotypic data on LR length from individual F_2_ plants at stage R6 (E1) and from adjusted progeny means for LR length at stage V6 (E2) were used. The F_3_ QTL mapping was performed on adjusted progeny means from F_3:4_ progenies averaged across trials (E3) for traits exhibiting significant genetic variance. Multiple QTL mapping was conducted using the “stepwiseqtl” function of the R/qtl package (Arends et al., 2010), which applies forward selection (up to 10 QTL) and backward elimination to identify the model with the highest penalized LOD score. Penalty values were derived from a 1000-iteration permutation test, with significance threshold of 0.01. For the most likely QTL model, we used the function “fitqtl” to obtain the proportion of phenotypic variance explained (PVE) for individual QTL and the full model, as well as for the estimation of additive and dominant QTL effects, by fitting all QTL simultaneously. Approximate 95% Bayesian credible QTL intervals were calculated with the function “bayesint”. The proportion of genetic variance explained was obtained by dividing the PVE by the trait heritability (Schön et al., 2004). For a given trait, the additive effects of the QTL were expressed in standard deviations by dividing the QTL additive effects by the standard deviation of the adjusted means of the respective trait. QTL were considered overlapping when their distance on the F_3_ genetic map was less than or equal to 20 cM. Detailed descriptions of the R/qtl functions used are provided in Broman and Sen (2009).

### qlr1 fine-mapping

In the fine-mapping experiments (E4-E7), 14 KASP markers on Chr. 1 (M1-M14) positioned within the 26 Mb region comprising *qlr1* were used. The statistical model to estimate the additive effect of M1-M14 in individual experiments E4-E7 and across E4-7 was:

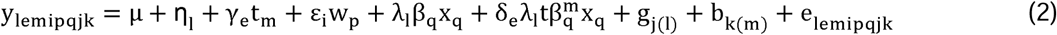

Where y_lemipqjk_ are the phenotypic observations, µ is the overall mean, η_l_ is the fixed effect of the group (l=1,2,3 for checks, RILs, HIFs), γ_e_ is a dummy variable with γ_e_ =1 in analyses comprising experiments with several trials (E4, E5, across E4-7) and γ_e_= 0 otherwise, t_m_ is the fixed effect of the trial m, ε_i_ is a dummy variable with ε_i_=1 for root traits assessed on separate whorls and ε_i_=0 otherwise, w_p_ is the fixed effect of the whorl p (p = 1 – 6 for whorls W2-W7), λ_l_ is a dummy variable with λ_l_=1 for HIFs and λ_l_=0 otherwise, β_q_ is the fixed additive effect of the tested marker q, x_q_ is the genotype score with x_q_ E [0,1,2] for genotypes AA-AB-BB (with AA being the genotype of P1, BB of P2 and AB the heterozygous genotype), δ_e_ is a dummy variable with δ_e_ =1 in analyses of experiments E4 and E5, each comprising two trials, and δ_e_= 0 otherwise, tβ^m^ is the fixed interaction of the additive effect of the tested marker q and the trial m, g_j(l)_ is the random effect of the genotype j nested in group l, b_k(m)_ is the random effect of block k nested in trial m, e_lemipqjk_ is the residual error. Random effects were assumed to be independent and identically normally distributed with mean zero and the respective variance component. The significance of the fixed effects was tested with an incremental Wald test implemented in ASReml-R. Fixed terms were considered significant for P < 0.05. The additive effects of the markers were expressed in standard deviations by dividing marker effects by the standard deviation of the adjusted means of the respective trait.

### Genomic resources and candidate gene analysis

Parental lines P1 and P2 were sequenced using PacBio HiFi long reads by CNRGV, INRAE Occitanie, Toulouse, France (cnrgv.toulouse.inrae.fr). Circular consensus sequences were de-novo assembled with hifiasm version 19 (Cheng et al., 2021) using default settings. Protein-coding gene sequences were detected using the MAKER toolset (Campbell et al., 2014), with cDNA and protein evidences based on other publicly available data. The genomic sequences of the annotated genes between M1 and M14 in the B73v5 reference genome were blasted against the assemblies of P1 and P2 to determine the respective genomic coordinates and obtain the underlying sequences. Multiple sequence alignment of the gene sequences of B73, P1 and P2, including all the gene sub-features (5’ untranslated regions (UTR), exons, coding sequences, 3’UTR), were performed with the “msa” package in R (Bodenhofer et al., 2015) and manually inspected for the genes within the fine-mapped region of *qlr1* (M4-M10). The genomic sequence of the annotated genes and respective sub-features for B73, their functional annotation, gene ontology terms, expression level in root tissues of B73 were retrieved from maize GDB (www.maizegdb.org) and refer to the latest annotation (Hufford et al., 2021).

### Transcript level measurements

In E7, five biological replicates of the distal 10 cm of crown roots of the last developed whorl at developmental stage R2 were harvested and flash-frozen in liquid nitrogen. From six genotypes (P1, P2, HIF6A, HIF6B, HIF7A, HIF7B) RNA was extracted using a guanidine hydrochloride protocol (Logemann et al., 1987), followed by DNase digestion (DNase I, RNase free, Thermo Scientific) and first-strand cDNA synthesis (Maxima H Minus Kit, random hexamer primers, Thermo Scientific K1652). RT-qPCRs were performed in three technical replicates with primers binding to the 3’UTR of the candidate gene Zm00001eb015500 (**Table S2**). The gene MEP (membrane protein PB1A10.07c, Zm00001eb257640) and LUG (Leunig, Zm00001eb357900) were used as references (Manoli et al., 2012). Normalization of RT-qPCR was performed with the delta-delta Cq method (Livak and Schmittgen, 2001).

## Results

### QTL for root and agronomic traits

In a preliminary field experiment (E0), we evaluated the adult root architecture of 18 doubled-haploid (DH) lines derived from the Austrian maize landrace “Kemater Landmais Gelb” at developmental stage R2. The two DH lines KE0413 (P1) and KE0113 (P2) differed significantly in lateral root (LR) length, representing the extremes for this trait, while showing similar flowering time and sharing a similar genomic background. By crossing P1 and P2 we developed the populations used in this study, which were evaluated in seven field and greenhouse experiments (**Figure 1**, **Table 1**).

The traits LR length, root dry weight (RDW), flowering time (FF), early and final plant height (PH_V6, PH_R2) and shoot dry weight (SDW) exhibited significant genetic variance in the bi-parental populations and served as the basis for QTL mapping analyses. LR length, assessed by visual scoring of the root stock (LRv) at stages V6, R2, R6 and with the software DIRT (Das et al., 2015) at stage R2 (LRd_R2) exhibited high heritability (h^2^) both in the DH lines (E0, h^2^=0.87) and in the populations derived from the bi-parental cross, with estimates of 0.74 for LRv_R2, 0.61 for LRd_R2, and 0.43 for LRv_V6 (**Table 1**).

QTL were detected for all traits except SDW_R2 (**Table 2**) and were distributed on all chromosomes, with some QTL co-localizing in the same genomic regions (**Figure 2**). For LR length, eight QTL were detected at R2, four at R6, and one QTL at V6 explaining 57%, 21%, and 13% of the phenotypic variance, respectively. The two QTL for LRd_R2 explained together 18% of the phenotypic variance. All four QTL for LR length at stage R6 co-localized with the respective QTL for LR length at stage R2 even though QTL mapping was performed in separate experiments (E1, E3). The QTL on Chr. 7 at stage V6 did not overlap with any of the QTL at reproductive stages. Despite a significant correlation of LR length assessed visually (LRv_R2) and with DIRT (LRd_R2) (r=0.58, P < 0.001, **Figure S3**), only one common QTL was found for the visual scoring and DIRT assessment (Chr. 1). These results suggest that the visual scoring on the root stock provided higher sensitivity than the measurement of LR length with DIRT from the excised root. The additive effects of the eight LRv_R2 QTL were moderate, ranging from 0.21 to 0.29 scores (corresponding to 0.25-0.35 standard deviations of the trait). For most LR length QTL, except for the QTL on Chr. 5 at 194 Mb (R2) and one QTL on Chr. 8 at 164 Mb, the trait increasing alleles were contributed by P1, explaining the large difference of LR length between parental lines. The LR length QTL distal on Chr. 9 (163 Mb) and Chr.10 (151 Mb) co-localized with QTL for flowering time, the latter being associated also with PH_V6. One QTL on Chr. 8 for LRv_R2 at 107 Mb co-localized with a QTL for root dry weight (RDW_R2), while all the other QTL for LR length were not associated with other traits. The QTL on Chr.1 overlapping for LRv_R2, LRv_R6, and LRd_R2 on Chr. 1, explaining the highest proportion of phenotypic variance for LR length at stage R2, was targeted for fine-mapping and will subsequently be referred to as *qlr1*.

**Table 2.** QTL identified for traits with significant genetic variance in experiments E1-E3. Given are the trait heritability (h^2^), chromosome (Chr) and physical position in Mb of the QTL [Pos (Mb)], estimated additive effect of the QTL P2 allele ± standard error (AE ± SE), proportion of phenotypic and genetic variance explained by the individual QTL [PVE (%), GVE (%)] and by the multi-QTL model [PVEm (%), GVEm (%)].

| Trait | $h^2$ | Chr | Pos (Mb) | AE ± SE | PVE (%) | PVEm (%) | GVE (%) | GVEm (%) |
| --- | --- | --- | --- | --- | --- | --- | --- | --- |
| LRv_R2 | 0.74 | 1 | 56 | -0.28 ± 0.04 | 9.2 | 57.4 | 12.4 | 77.6 |
| | | 3 | 187 | -0.29 ± 0.04 | 8.9 | | 12.0 | |
| | | 5 | 6 | -0.25 ± 0.04 | 6.4 | | 8.7 | |
| | | 5 | 194 | 0.24 ± 0.04 | 7.3 | | 9.8 | |
| | | 8 | 20 | -0.23 ± 0.04 | 5.1 | | 6.8 | |
| | | 8 | 107 | -0.25 ± 0.04 | 5.5 | | 7.4 | |
| | | 8 | 164 | 0.23 ± 0.04 | 6.4 | | 8.6 | |
| | | 9 | 163 | -0.21 ± 0.04 | 4.6 | | 6.3 | |
| LRv_R6 | | 1 | 52 | -0.49 ± 0.08 | 5.8 | 21.2 | | |
| | | 5 | 13 | -0.39 ± 0.09 | 4.2 | | | |
| | | 5 | 206 | 0.61 ± 0.09 | 8.0 | | | |
| | | 8 | 17 | -0.43 ± 0.08 | 3.7 | | | |
| LRv_V6 | 0.43 | 7 | 19 | -0.31 ± 0.07 | 13.3 | 13.3 | 30.8 | 30.8 |
| LRd_R2 | 0.61 | 1 | 54 | -1.76 ± 0.29 | 10.9 | 18.5 | 17.8 | 30.3 |
| | | 10 | 151 | -1.41 ± 0.31 | 7.9 | | 13.0 | |
| RDW_R2 | 0.65 | 6 | 176 | -2.99 ± 0.55 | 9.4 | 20.4 | 14.4 | 31.5 |
| | | 8 | 109 | -3.15 ± 0.53 | 10.3 | | 15.9 | |
| FF | 0.88 | 1 | 22 | -0.87 ± 0.13 | 6.9 | 58.5 | 7.9 | 66.5 |
| | | 2 | 3 | 0.80 ± 0.14 | 6.2 | | 7.0 | |
| | | 3 | 5 | 0.68 ± 0.13 | 4.2 | | 4.8 | |
| | | 7 | 2 | 0.64 ± 0.13 | 3.9 | | 4.5 | |
| | | 7 | 185 | -0.74 ± 0.13 | 4.9 | | 5.6 | |
| | | 9 | 160 | -1.18 ± 0.13 | 12.1 | | 13.7 | |
| | | 10 | 151 | 1.31 ± 0.13 | 18.3 | | 20.8 | |
| PH_V6 | 0.75 | 10 | 151 | -3.93 ± 0.38 | 31.2 | 31.2 | 41.6 | 41.6 |
| PH_R2 | 0.71 | 6 | 158 | 3.51 ± 0.59 | 10.9 | 19.4 | 15.3 | 27.3 |
| | | 8 | 142 | -3.37 ± 0.58 | 10.2 | | 14.3 | |

Trait codes: LRv, visual scoring of LR length (at stages V6, R2, R6); LRd, DIRT assessment of LR length (at stage R2), FF, flowering time; PH plant height (at stages V6, R2), RDW root dry weight (at stage R2).

### Assessment of LR length in different whorls

In the fine-mapping experiments E4-E7 we dissected the root system and measured LR length on separate whorls (W), from W2 to W7. The shoot-borne roots accessible at stage R2 capture the root system’s developmental history, with older whorls being still accessible and no new ones emerging, as the root system reached full maturity. Analogously to our findings on the root stock (RS, **Figure S3**), LR length measured with DIRT (LRd) on whorl-specific excised roots correlated strongly with the visual scoring (LRv) (**Figure S4**), with the latter displaying slightly higher heritability (**Figure S5**). Consequently, LRv was used for assessing LR length both for RS and excised roots of individual whorls. Across trials and genotypes, LR length increased from W2 to W5, decreased from W5 to W7, and showed intermediate to high positive correlations between whorls, suggesting shared genetic mechanisms and negligible intra-whorl competition (**Figure 3**). LR length assessed from the RS was positively correlated with LR length scored on separate whorls (particularly with the later forming whorls W4, W5, and W6), and gave the highest heritability (h^2^ = 0.75 across E4-7). Thus, the visual scoring on the RS could be shown to be a fast and comprehensive measure of LR length for the mature root system.

**Figure 3.**
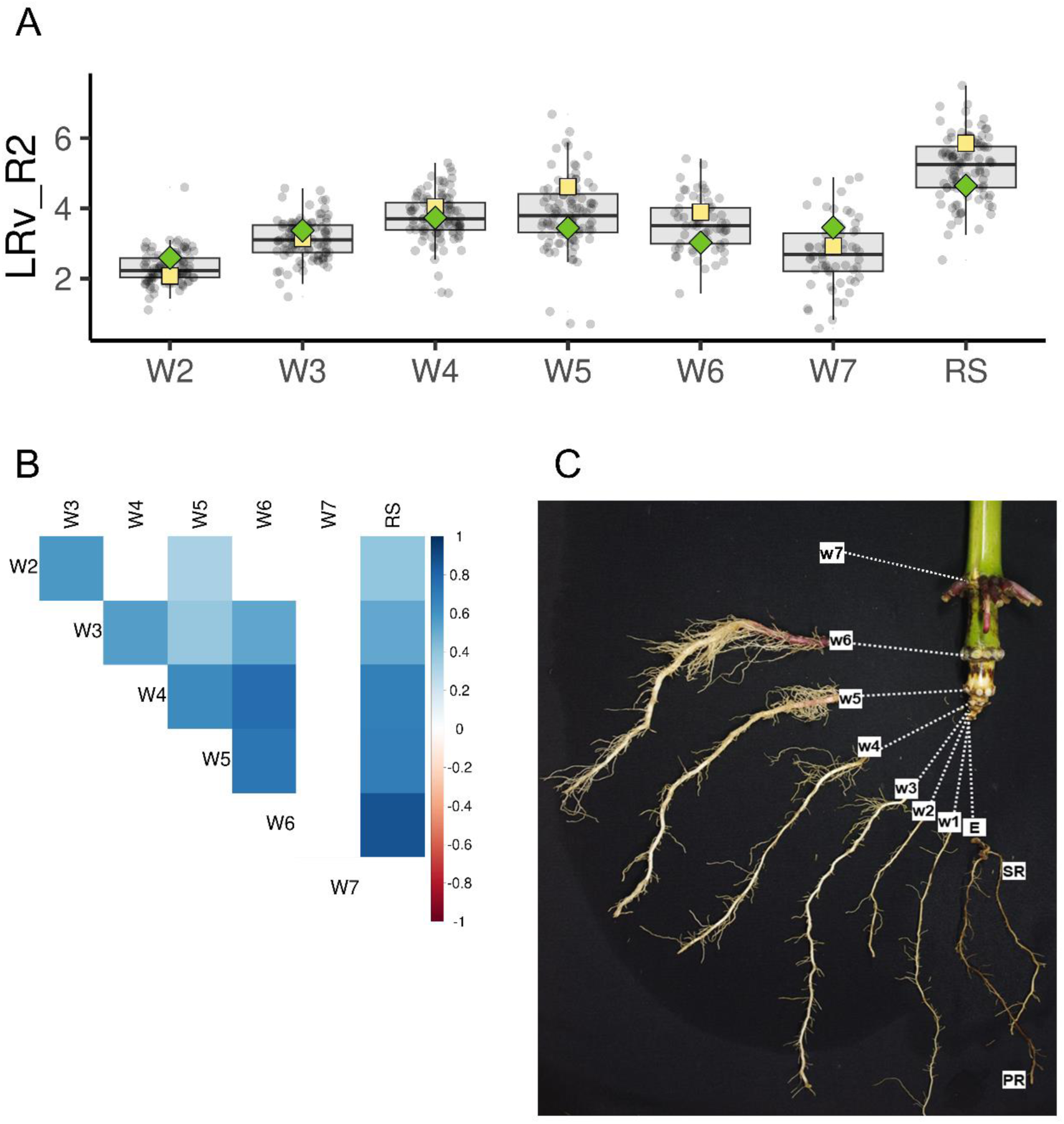
Assessment of lateral root (LR) length in different whorls by visual scoring at developmental stage R2 (LRv_R2). **A)** Distributions of LRv_R2 measured on the root stock (RS) and excised roots of whorls W2-W7 of 91 HIF progenies, 4 RILs and 5 checks. Dots represent adjusted means across experiments (E4-7, n=100). P1 and P2 are shown as yellow squares and green diamonds, respectively. Boxplots show medians (thick lines), first and third quartiles (hinges), and whiskers extending to 1.5 times the interquartile range. **B)** Pairwise Pearson correlations of LR length measured from the RS and W2-W7. Only significant correlations (P < 0.05, Bonferroni–Holm corrected) are displayed. **C)** Root system dissection of a representative plant from this study at stage R2. Embryonic roots (E), including the primary root (PR) and seminal roots (SR), and shoot-borne roots of whorls W1-W7 are visible.

### Fine-mapping of qlr1

In 43 heterogeneous inbred families (HIFs) with similar genomic background and segregating for the *qlr1* QTL, marker-trait associations detected for LR length across experiments E4-7 pointed consistently to the region between markers M5 (AX-90529220, pos. 51,364,010 bp) and M9 (AX-90529237, pos. 52,988,306 bp) for both RS and averaged across whorls (**Figure 4A**).

**Figure 4.**
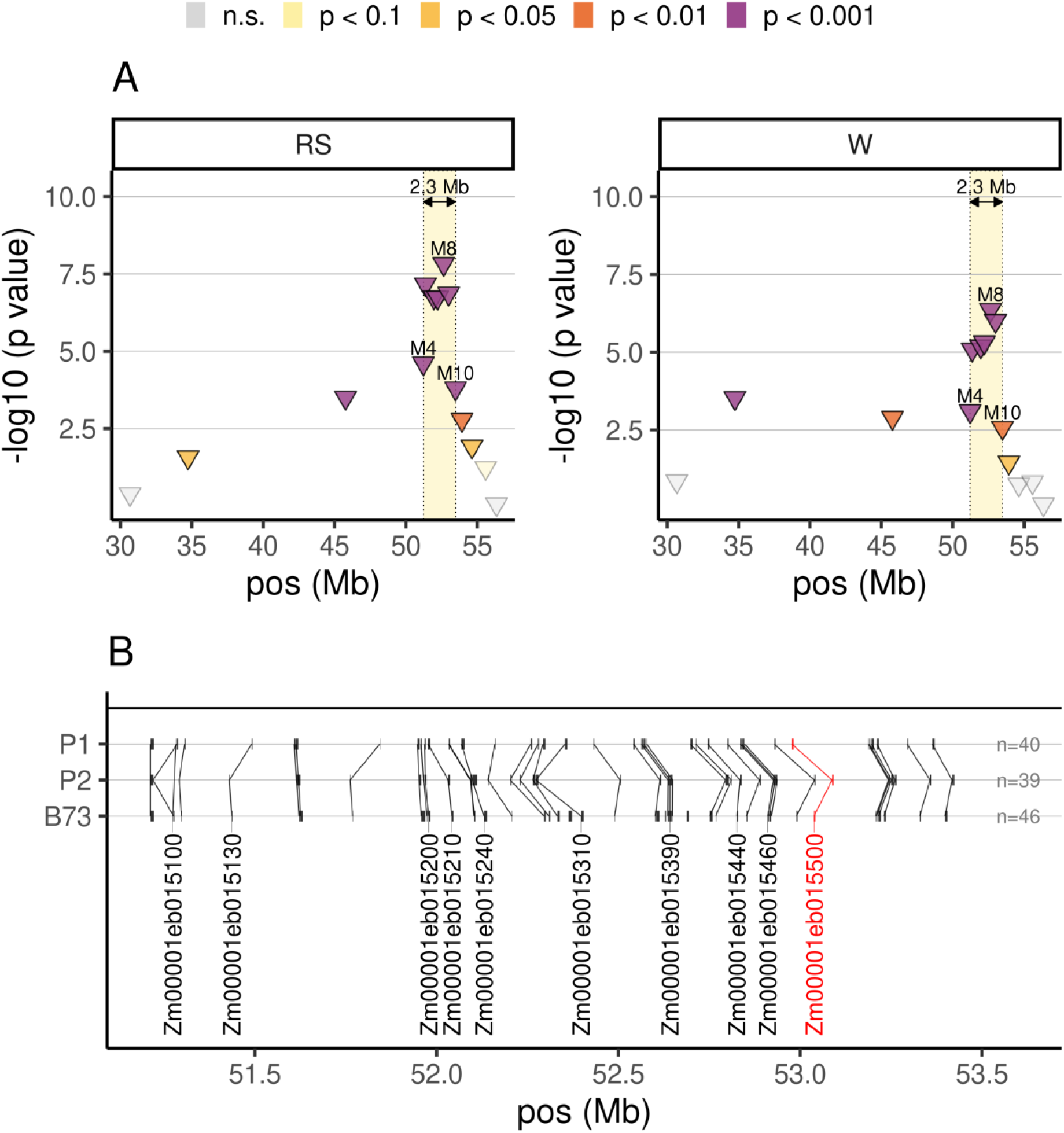
Fine mapping of QTL *qlr1*. **A)** Significance of the marker-trait associations of 14 markers located within *qlr1* with lateral root (LR) length assessed by visual scoring at developmental stage R2 (LRv_R2) across experiments (E4-7). LRv_R2 was measured on the RS and averaged across excised roots from W2-W7 (W). Markers are represented by triangles and coloured according to the level of significance of the marker-trait associations. The fine-mapped region (M4–M10) is indicated by a dotted box shaded in yellow. Flanking markers M4 and M10 and the most significant marker M8 are labelled. The X-axis represents the physical position on chromosome 1 based on B73v5 in Megabases (Mb). **B)** Annotated genes between M4 and M10 in P1 (KE0413), P2 (KE0113) and B73 (B73v5). Genes are represented as vertical segments in the respective genomic positions and connected if syntenic. For visualization on a common axis, the starting coordinate of the first gene of B73 is taken as reference for all genomes. Ten prioritized candidate genes are labelled. Zm00001eb015500, the candidate of highest priority, is labelled in red.

Significance of marker-trait associations dropped sharply for flanking markers M4 (AX.91366105, pos. 51,218,801 bp) and M10 (AX.91334117, pos. 53,476,179), thus identifying a 2.3 Mb interval on Chr. 1 delimited by M4 and M10 to harbor candidate genes for LR length. The 2.3 Mb region contained 46 protein-coding gene models annotated in B73v5, of which 40 and 39 were syntenic in P1 and P2, respectively (**Figure 4B**).

A prioritized subset of 10 candidate genes was identified based on their functional annotations, literature search, expression in B73 root tissues, and polymorphisms between P1 and P2 (**Table S3**). Among these, Zm00001eb015500 (hb130), a transcription factor functionally annotated for adventitious root development and LR formation, emerged as a potential candidate. P1 and P2 alleles of hb130 differed for a tandem insertion in the first intron and several polymorphisms resulting in two amino acid substitutions and a small insertion in the protein product. In E7, RT-qPCR revealed significantly higher transcript abundance of the P2 as compared to the P1 allele in parental lines and two HIFs selected to segregate for the hb130 allele (P < 0.01), indicating that these alleles differ in their gene expression levels (**Figure S6**).

### qlr1 characterization

Marker M8 (AX-90529242, pos. 52643392) showed consistently the highest significance across experiments and was selected as representative marker for *qlr1*. In the 43 HIFs, the P1 allele of M8 was significantly associated with increased LR length both when assessed on the RS and on individual whorls (**Figure 5A**). Across E4-7, the *qlr1* effect on LR length was significant in all whorls except for W3, and most pronounced in later forming whorls W5 and W6. The *qlr1* additive effect of 0.23-0.25 scores for LR length on the RS, W5 and W6 was in line with the results from the QTL mapping experiment (E3). *qlr1* impacted LR length significantly in all experiments (E4-E7), albeit effect sizes varied across environments and trials (**Figure 5B**). Significant marker associations (P < 0.001) were also detected for LR length assessed with DIRT on excised roots averaged across whorls (LRd_R2w) and plant height at stages V6 and R2 (PH_V6, PH_R2), with the P1 allele increasing trait expression in all cases (**Figure 5C**).

**Figure 5.**
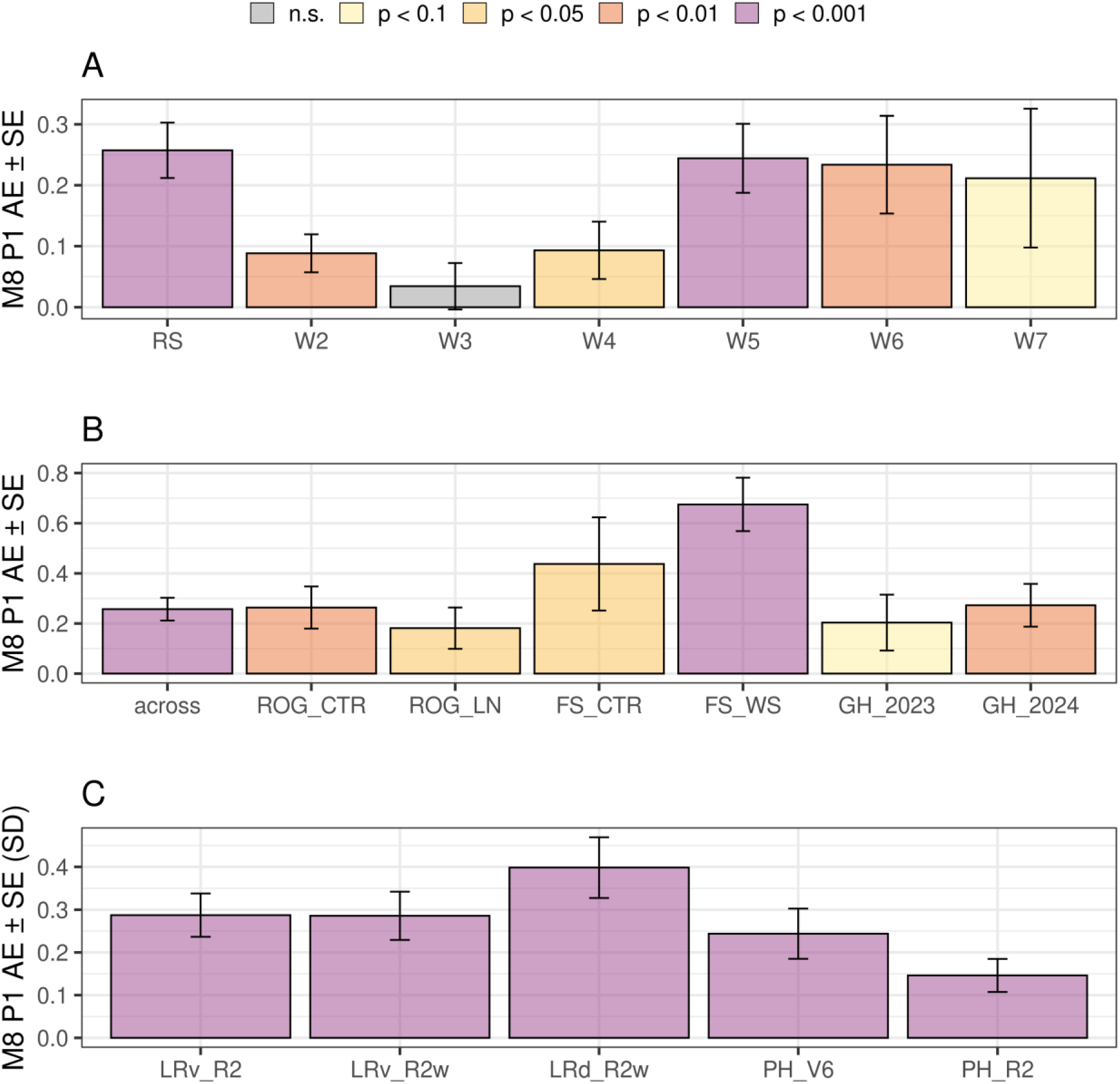
*qlr1* characterization. Additive effects of the P1 allele at QTL *qlr1* estimated based on the association of marker M8 with lateral root (LR) length and plant height. LR length was assessed by visual scoring at developmental stage R2 (LRv_R2) in experiments E4-E7. **A)** Allele effects for LRv_R2 measured on the root stock (RS) and separate whorls (W2-W7) on 43 HIFs across experiments (E4-7). **B)** Allele effects for LRv_R2 on the RS across experiments E4-7 and in individual trials, estimated from 43 HIFs across experiments; 25 HIFs in ROG, 3 HIFs in FS, 13 HIFs in GH_2023 and 9 HIFs in GH_2024. **C)** Allele effects for different LR traits and plant height at stages V6 and R2 significantly associated with marker M8 across experiments E4-7, expressed as standard deviations (SD) of the trait. Bars represent the additive effect ± standard error (AE ± SE) estimated at marker M8 with colours indicating differences in the level of significance of the marker-trait associations. **Trial abbreviations**: ROG, Roggenstein; FS, Freising; CTR, control; LN, low N; WS, water stress; GH, Greenhouse; 2023, 2024 year. **Trait abbreviations**: LRv_R2w, LR length across whorls (visual scoring); LRd_R2w, LR length across whorls (DIRT); PH_V6, PH_R2, plant height (stages V6 and R2).

### Correlating shoot weight and LR length under different N, P, and water treatments

In E3-5, we evaluated 289 RILs and 25 HIFs segregating for *qlr1*, flint maize inbred lines, and the parents of the mapping populations under control conditions and limited phosphorus (P), nitrogen (N) and water (W) availability, detecting significant root and shoot responses to these treatments. Effects varied for different traits and developmental stages (**Table S4**).

Here we focus on the impact of these treatments on LR length (LRv_R2) and shoot dry weight (SDW_R2), as well as their correlation. SDW_R2 served as a proxy for plant productivity, reflecting the cumulative biomass accumulation over the growing period until stage R2.

Under N limited conditions (E4), both LR length and SDW_R2 were significantly reduced (P < 0.001) compared to the control treatment (**Figure 6A**). The two traits were significantly correlated under control conditions (r = 0.29, P < 0.05), but not under N stress (**Figure 6B**), suggesting that reduced N availability disrupted the association between the two traits. The P limited conditions (E3) had neither an effect on LR length nor on shoot weight compared to the control (**Figure 6C**) and correlations between the two traits were significant but small in both treatments (r ≤ 0.30) (**Figure 6D**).

**Figure 6.**
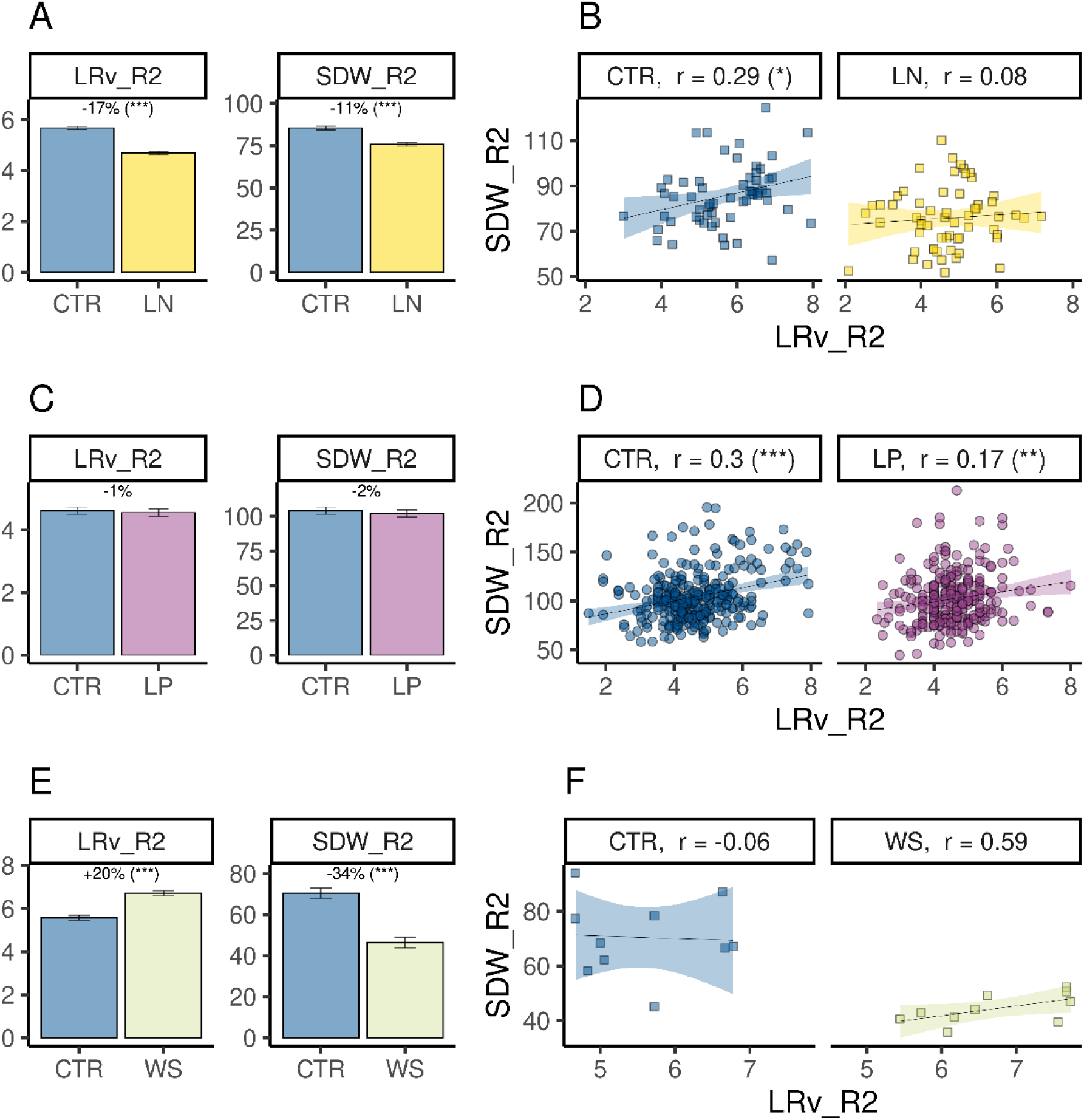
Functional consequences of lateral root (LR) length. Effect of nutrient and water treatments on LR length assessed by visual scoring (LRv_R2) and shoot dry weight (SDW_R2) as well as their correlation. **A, C, E)** Average LRv_R2 and SDW_R2 under control and treatment conditions. Bars represent the adjusted treatment means ± standard error. The treatment effect, expressed as percentage relative to control, along with its significance, is indicated above the bars. **B, D,F)** Scatterplot of LR length (X-axis) and SDW (Y-axis). Dots represent adjusted means within trials. The Pearson correlation coefficient and its significance are shown above each scatterplot. The regression line and its associated confidence interval (shaded area) were fitted with the function “geom_smooth” in ggplot2. Underlying data came from experiment E4 (A, B), E3 (C, D) and E5 (E, F). For data dimensions, see Table 1. Significant differences are marked with stars: *** P < 0.001, ** P < 0.01, * P < 0.05. **Treatment abbreviations**: CTR, control; LP, low phosphorous; LN, low nitrogen; WS, water stress.

Under limited water supply (E5), LR length was significantly increased compared to control conditions (P < 0.001), while SDW_R2 was significantly reduced (P < 0.001) (**Figure 6E**). No significant correlation could be observed between the two traits in either of the two treatments, most likely due to the small sample size (n=10) (**Figure 6F**).

## Discussion

Adapting the root architecture to target environments can enhance production stability and sustainability of our crops. Breeding maize varieties with this goal in mind necessitates a comprehensive understanding of the genetic regulation of root traits and their functional consequences on yield, yield stability, and resource use efficiency. Among root traits, lateral root (LR) architecture is of prime importance (Lynch, 2018, 2022). In this study, we leveraged the native diversity of an Austrian maize landrace to understand the genetic basis of LR length and other root and shoot traits across the life cycle of the maize plant.

Our knowledge on the genetic control of LR development on adult, shoot-borne roots in maize is limited also due to the lack of efficient phenotyping methods. This limitation makes it practically impossible to evaluate large numbers of plants, a key prerequisite in plant breeding. Here we showed that the visual scoring of LR length from root stock (RS) pictures provided a fast and robust phenotyping method to quantify LR length with high heritability and a strong correlation with the more time-consuming assessment from excised shoot-borne roots.

Both in the landrace-derived doubled-haploid (DH) lines and in the populations derived from a bi-parental cross, estimates of LR length heritability exceeded previously reported values (Burton et al., 2014; Schneider et al., 2020), indicating a substantial genetic component contributing to trait expression in the landrace-derived plant material. This allowed us to identify four QTL for LR length with high consistency across experiments, developmental stages, and environments. The overlap of QTL at reproductive stages R2 and R6 and their absence at stage V6 suggests that distinct genes may have influenced LR elongation of shoot-borne roots at different stages, while LR architecture stabilized post-R2 and was preserved until maturity (R6), despite senescence and root degradation. However, this warrants further research as differences in plant development at harvest, the smaller population size and the greenhouse growing conditions in E2 might have contributed to these differences in the vegetative and the reproductive stages. All the QTL we identified for LR length had moderate to small effect sizes, consistent with a quantitative genetic architecture, similarly to what was reported for other shoot-borne root QTL (Burton et al., 2014; Zhang et al., 2018c; Ren et al., 2022; Li et al., 2024).

Mapping a QTL at high genetic resolution helps to avoid linkage drag when the QTL is introgressed by marker-based breeding and provides the starting point for candidate gene identification and map-based cloning. Dissecting the root system at stage R2 allowed us to measure root traits from shoot-borne roots formed over time by analysing individual whorls and assess the relationships and contributions of individual whorls to overall root architecture. The QTL *qlr1* consistently affected LR length of the root stock and of excised roots of different whorls in different growing conditions and nutrient regimes and across different environments. This suggests that the *qlr1* locus contributes to the constitutive alteration of the root system by modulating the LR length of shoot-borne roots.

The stable effect of *qlr1* in combination with the large number of heterogeneous inbred families (HIFs) tested in E4-E7, allowed us to fine-map *qlr1* to a 2.3 Mb region comprising 46 genes annotated in B73v5. Based on whole genome sequence information of the two parents P1 and P2 and genomic data from publicly available datasets from maize and other species, we identified a possible candidate gene in the fine-mapped *qlr1* interval influencing LR length and possibly other root and shoot traits: hb130, a transcription factor with homeobox domain. Homologous genes to hb130 have been shown to modify root and plant architecture in rice (*Oryza sativa*) and Arabidopsis (*Arabidopsis thaliana*), including LR formation (Sheng et al., 2017), LR primordium size (Kawai et al., 2022), crown root (Zhao et al., 2009; Cheng et al., 2016; Zhou et al., 2017; Zhang et al., 2018b) and tiller formation (Zhang et al., 2018a). Albeit primarily expressed in the embryo (Chen et al., 2024), its expression in crown roots observed in our study aligned with the LR length phenotype. The differential expression of hb130 in parents P1 and P2 and in HIFs is possibly linked to the tandem insertion in the first intron of the allele of P1, which is the most likely variant impacting transcription rate and/or post-transcriptional regulation, as the gene sequence and core promoter elements are otherwise highly similar in P1 and P2. The role of hb130 homologs in regulating root architecture appears to be context-dependent, as they are part of complex regulatory networks responsive to both genetic and environmental cues. For example, overexpression of wox11, the closest Arabidopsis ortholog of hb130, reduced LR density in plants grown *in vitro*. Interestingly, also the repression of the same gene led to reduced LR density, but only in soil-grown plants, suggesting environment-specific gene function (Sheng et al., 2017). In maize, knockout of hb77, an ancient paralog of hb130, decreased seminal root number while increasing LR density, suggesting pleiotropic gene action (Yu et al., 2024). Thus, establishing a direct link between reduced hb130 expression and increased LR length requires additional evidence, which is beyond the scope of this study and warrants future research. At the current stage, we cannot exclude that other genes in the region may also contribute to the effect of QTL *qlr1*.

To evaluate the functional impact of the LR length phenotype, we analysed biomass accumulation of RILs and HIFs varying for LR length under control conditions and limited nitrogen (N) and phosphorus (P) availability. We observed that genotypes with longer LR tended to accumulate more shoot biomass in control conditions, while this trend was absent under low N. Under low P, genotypes experienced a reduction of early vigour and root biomass, but not of shoot biomass. Therefore, we cannot exclude that, despite the naturally low P content in the experimental field, its availability was not sufficiently limiting to constrain plant growth and be considered a stress factor. The relationship we observed between LR length and shoot biomass accumulation under varying N and P availability was weaker than previously reported (Zhan and Lynch, 2015; Jia et al., 2018), and genotypes with long LR performed similar or better than genotypes with short LR in all tested conditions. We therefore conclude that, in the absence of trade-offs, increased LR length may confer a functional advantage for the plant by increasing resource uptake and thus supporting shoot biomass accumulation.

In this study, we showed that several QTL with small to moderate effects influenced LR length of shoot-borne roots at different developmental stages in maize, we characterized and fine-mapped the most significant of these QTL, *qlr1*, to a 2.3 Mb region, suggested a potential candidate gene, and assessed the functional consequences of LR length under different growing conditions, finding a positive correlation with biomass accumulation in control conditions which was disrupted under N stress. The extensive genetic diversity pertaining to landraces represents a valuable resource to promote genetic studies on root traits. Examples of landrace alleles used for modifying root architecture and enhancing soil resource uptake exist in rice (Gamuyao et al., 2012; Uga et al., 2013). Before introgressing the allele for increased LR length into elite material, its interaction with the genetic background of the recipient population needs to be investigated. Marker-based selection can facilitate this task as well as the identification of additional allelic variation at the other QTL we identified for LR length. Using *qlr1* flanking markers and additional marker developments within the 2.3 Mb target, fine-mapping & map-based cloning of the causal feature can be achieved. In addition, exploring natural allelic variation or generating new variation (via chemical induced mutations, Genome Editing, or classical transgenic approaches) at the *qlr1* locus in breeding material may allow the modification of LR length in adult shoot-borne roots and facilitate the breeding of elite genotypes with modified nutrient and water uptake to suit specific environmental conditions.

## Supplementary data

Table S1. List of traits assessed in experiments E0-E7 and their definitions.

Table S2. Primer pairs used to measure the transcript levels of hb130.

Table S3. Information on candidate genes in the fine mapped region of the QTL *qlr1*.

Table S4. Effect of nitrogen, phosphorous and irrigation treatments on root and agronomic traits.

Figure S1. Scoreboards for visual scoring of LR length.

Figure S2. Genetic maps used for QTL mapping projected onto the B73v5 physical map. Figure S3. Correlation of LR length assessed on the root stock with visual scoring and DIRT. Figure S4. Correlation of LR length assessed on excised roots with visual scoring and DIRT. Figure S5. Heritability of LR length assessed on excised roots with visual scoring and DIRT. Figure S6. Characterization of candidate gene Zm00001eb015500 (hb130).

## Acknowledgments

We thank Brigitte Neuhauser, Iris Prücklmaier, Sylwia Schepella, Stefan Schwertfirm, Margot Siebler, Stefan Kimmelmann, Sonja Biedefeld, Tanja Rettig, and Nadine Gautier for technical assistance. We thank Prof. Dr. Peter Westhoff and Dr. Monika Frey for valuable insights and constructive suggestions that contributed to improve this manuscript. We thank the Plant Technology Center (Technical University of Munich, Germany) for providing infrastructure and technical support during greenhouse and field experiments. We acknowledge the GENTYANE platform of Clermont-Ferrand INRAE Center (https://gentyane.clermont.inrae.fr/) for providing assistance in NGS sequencing.

## Author contributions

F.G., C-C.S., and M.O. designed the research and developed ideas; M.O., T.P., D.S., C.U and F.G. developed and genotyped the plant material; F.G, T.P., and S.S. designed and performed phenotyping experiments, C.C generated the whole genome sequencing data, F.G. analyzed phenotypic and genotypic data, F.G, S.R and C.C. analyzed the whole genome sequencing data; F.G., S.S., and C.-C.S. wrote the manuscript; all authors read and approved the final manuscript; C.-C.S. agrees to serve as the author responsible for contact and to ensure communication.

## Data availability

All primary data to support the findings of this study are available upon request to the corresponding author.

## References

1. Ahmed MA, Zarebanadkouki M, Meunier F, Javaux M, Kaestner A, Carminati A. 2018. Root type matters: measurement of water uptake by seminal, crown, and lateral roots in maize. Journal of Experimental Botany 69: 1199–1206

2. Arends D, Prins P, Jansen RC, Broman KW. 2010. R/qtl: high-throughput multiple QTL mapping. Bioinformatics 26: 2990–2992

3. Baer M, Taramino G, Multani D, Sakai H, Jiao SP, Fengler K, Hochholdinger F. 2023. Maize *lateral rootless 1* encodes a homolog of the DCAF protein subunit of the CUL4-based E3 ubiquitin ligase complex. New Phytologist 237: 1204–1214

4. Bodenhofer U, Bonatesta E, Horejs-Kainrath C, Hochreiter S. 2015. msa: an R package for multiple sequence alignment. Bioinformatics 31: 3997–3999

5. Broman KW, Sen S. 2009. A Guide to QTL Mapping with R/qtl. Springer

6. Burton AL, Johnson JM, Foerster JM, Hirsch CN, Buell CR, Hanlon MT, Kaeppler SM, Brown KM, Lynch JP. 2014. QTL mapping and phenotypic variation for root architectural traits in maize (*Zea mays* L.). Theoretical and Applied Genetics 127: 2293–2311

7. Butler DG, Cullis, B.R.A., R. Gilmour, Gogel, B.G. and Thompson, R. 2017. ASReml-R Reference Manual Version 4. VSN International Ltd, Hemel Hempstead, HP1 1ES, UK

8. Campbell MS, Holt C, Moore B, Yandell M. 2014. Genome annotation and curation using MAKER and MAKER-P. Current protocols in bioinformatics 48: 4.11. 11-14.11. 39

9. Chen XX, Hou YY, Cao YY, Wei B, Gu L. 2024. A Comprehensive Identification and Expression Analysis of the WUSCHEL Homeobox-Containing Protein Family Reveals Their Special Role in Development and Abiotic Stress Response in *Zea mays* L. International Journal of Molecular Sciences 25

10. Cheng HY, Concepcion GT, Feng XW, Zhang HW, Li H. 2021. Haplotype-resolved de novo assembly using phased assembly graphs with hifiasm. Nature Methods 18: 170–175

11. Cheng SF, Zhou DX, Zhao Y. 2016. *WUSCHEL*-related homeobox gene *WOX11* increases rice drought resistance by controlling root hair formation and root system development. Plant Signaling & Behavior 11

12. Das A, Schneider H, Burridge J, Ascanio AKM, Wojciechowski T, Topp CN, Lynch JP, Weitz JS, Bucksch A. 2015. Digital imaging of root traits (DIRT): a high-throughput computing and collaboration platform for field-based root phenomics. Plant Methods 11

13. Gamuyao R, Chin JH, Pariasca-Tanaka J, Pesaresi P, Catausan S, Dalid C, Slamet-Loedin I, Tecson-Mendoza EM, Wissuwa M, Heuer S. 2012. The protein kinase Pstol1 from traditional rice confers tolerance of phosphorus deficiency. Nature 488: 535–539

14. Gautam V, Singh A, Yadav S, Singh S, Kumar P, Das SS, Sarkar AK. 2021. Conserved *LBL1-ta-siRNA* and miR165/166-*RLD1/2* modules regulate root development in maize. Development 148

15. Guffanti F, Nagel K, Pariyar S, Scheuermann D, Urbany C, Presterl T, Ouzunova M, Schoen CC. unpublished. GWAS reveals QTL for root traits in maize landraces.

16. Hochholdinger F, Park WJ, Sauer M, Woll K. 2004. From weeds to crops: genetic analysis of root development in cereals. Trends in Plant Science 9: 42–48

17. Hochholdinger F, Yu P, Marcon C. 2018. Genetic Control of Root System Development in Maize. Trends in Plant Science 23: 79–88

18. Hoelker AC, Mayer M, Presterl T, Bolduan T, Bauer E, Ordas B, Brauner PC, Ouzunova M, Melchinger AE, Schoen CC. 2019. European maize landraces made accessible for plant breeding and genome-based studies. Theoretical and Applied Genetics 132: 3333–3345

19. Holland JB, Nyquist WE, Cervantes-Martínez CT, Janick J. 2003. Estimating and interpreting heritability for plant breeding: an update. Plant breeding reviews 22

20. Holm S. 1979. A simple sequentially rejective multiple test procedure. Scandinavian journal of statistics: 65–70

21. Hufford MB, Seetharam AS, Woodhouse MR, et al. 2021. De novo assembly, annotation, and comparative analysis of 26 diverse maize genomes. Science 373: 655–662

22. Jia XC, Liu P, Lynch JP. 2018. Greater lateral root branching density in maize improves phosphorus acquisition from low phosphorus soil. Journal of Experimental Botany 69: 4961–4970

23. Kawai T, Shibata K, Akahoshi R, et al. 2022. *WUSCHEL-related homeobox* family genes in rice control lateral root primordium size. Proceedings of the National Academy of Sciences of the United States of America 119

24. Li PC, Zhang ZH, Xiao G, et al. 2024. Genomic basis determining root system architecture in maize. Theoretical and Applied Genetics 137

25. Livak KJ, Schmittgen TD. 2001. Analysis of relative gene expression data using real-time quantitative PCR and the 2− ΔΔCT method. methods 25: 402–408

26. Logemann J, Schell J, Willmitzer L. 1987. IMPROVED METHOD FOR THE ISOLATION OF RNA FROM PLANT-TISSUES. Analytical Biochemistry 163: 16–20

27. Lu CC, Chen MX, Liu R, et al. 2019. Abscisic Acid Regulates Auxin Distribution to Mediate Maize Lateral Root Development Under Salt Stress. Frontiers in Plant Science 10

28. Lynch JP. 2007. Roots of the second green revolution. Australian Journal of Botany 55: 493–512

29. Lynch JP. 2018. Rightsizing root phenotypes for drought resistance. Journal of Experimental Botany 69: 3279–3292

30. Lynch JP. 2022. Harnessing root architecture to address global challenges. Plant Journal 109: 415–431

31. Manoli A, Sturaro A, Trevisan S, Quaggiotti S, Nonis A. 2012. Evaluation of candidate reference genes for qPCR in maize. Journal of plant physiology 169: 807–815

32. Mayer M, Hölker AC, González-Segovia E, Bauer E, Presterl T, Ouzunova M, Melchinger AE, Schön CC. 2020. Discovery of beneficial haplotypes for complex traits in maize landraces. Nature Communications 11

33. Mayer M, Hölker AC, Presterl T, Ouzunova M, Melchinger AE, Schön C-C. 2022. Genetic diversity of European maize landraces: Dataset on the molecular and phenotypic variation of derived doubled-haploid populations. Data in Brief 42: 108164

34. Mayer M, Unterseer S, Bauer E, de Leon N, Ordas B, Schön CC. 2017. Is there an optimum level of diversity in utilization of genetic resources? Theoretical and Applied Genetics 130: 2283–2295

35. Mueller ND, Gerber JS, Johnston M, Ray DK, Ramankutty N, Foley JA. 2012. Closing yield gaps through nutrient and water management. Nature 490: 254–257

36. Peng YF, Li XX, Li CJ. 2012. Temporal and Spatial Profiling of Root Growth Revealed Novel Response of Maize Roots under Various Nitrogen Supplies in the Field. Plos One 7

37. Peng YF, Niu JF, Peng ZP, Zhang FS, Li CJ. 2010. Shoot growth potential drives N uptake in maize plants and correlates with root growth in the soil. Field Crops Research 115: 85–93

38. R Core Team. 2025. R: A Language and Environment for Statistical Computing. *In*, Ed 4.4.3. R Foundation for Statistical Computing

39. Ren W, Zhao LF, Liang JX, et al. 2022. Genome-wide dissection of changes in maize root system architecture during modern breeding. Nature Plants 8: 1408–1422

40. Schneider HM, Klein SP, Hanlon MT, Nord EA, Kaeppler S, Brown KM, Warry A, Bhosale R, Lynch JP. 2020. Genetic control of root architectural plasticity in maize. Journal of Experimental Botany 71: 3185–3197

41. Schön CC, Utz HF, Groh S, Truberg B, Openshaw S, Melchinger AE. 2004. Quantitative trait locus mapping based on resampling in a vast maize testcross experiment and its relevance to quantitative genetics for complex traits. Genetics 167: 485–498

42. Sheng LH, Hu XM, Du YJ, Zhang GF, Huang H, Scheres B, Xu L. 2017. Non-canonical *WOX11*-mediated root branching contributes to plasticity in *Arabidopsis* root system architecture. Development 144: 3126–3133

43. Singh V, van Oosterom EJ, Jordan DR, Messina CD, Cooper M, Hammer GL. 2010. Morphological and architectural development of root systems in sorghum and maize. Plant and Soil 333: 287–299

44. Trachsel S, Kaeppler SM, Brown KM, Lynch JP. 2011. Shovelomics: high throughput phenotyping of maize (Zea mays L.) root architecture in the field. Plant and Soil 341: 75–87

45. Uga Y, Sugimoto K, Ogawa S, et al. 2013. Control of root system architecture by *DEEPER ROOTING 1* increases rice yield under drought conditions. Nature Genetics 45: 1097–1102

46. Unterseer S, Bauer E, Haberer G, et al. 2014. A powerful tool for genome analysis in maize: development and evaluation of the high density 600 k SNP genotyping array. Bmc Genomics 15

47. Urzinger S, Avramova V, Frey M, Urbany C, Scheuermann D, Presterl T, Reuscher S, Ernst K, Mayer M, Marcon C. 2025. Embracing native diversity to enhance the maximum quantum efficiency of photosystem II in maize. Plant Physiology 197: kiae670

48. Viana WG, Scharwies JD, Dinneny JR. 2022. Deconstructing the root system of grasses through an exploration of development, anatomy and function. Plant Cell and Environment 45: 602–619

49. von Behrens I, Komatsu M, Zhang YX, Berendzen KW, Niu XM, Sakai H, Taramino G, Hochholdinger F. 2011. *Rootless with undetectable meristem 1* encodes a monocot-specific AUX/IAA protein that controls embryonic seminal and post-embryonic lateral root initiation in maize. Plant Journal 66: 341–353

50. Wang HM, Tang X, Yang XY, Fan YY, Xu Y, Li PC, Xu CW, Yang ZF. 2021. Exploiting natural variation in crown root traits via genome-wide association studies in maize. Bmc Plant Biology 21

51. Woll K, Borsuk LA, Stransky H, Nettleton D, Schnable PS, Hochholdinger F. 2005. Isolation, characterization, and pericycle-specific transcriptome analyses of the novel maize lateral and seminal root initiation mutant *rum1*. Plant Physiology 139: 1255–1267

52. Yu P, Gutjahr C, Li CJ, Hochholdinger F. 2016. Genetic Control of Lateral Root Formation in Cereals. Trends in Plant Science 21: 951–961

53. Yu P, Li CH, Li M, et al. 2024. Seedling root system adaptation to water availability during maize domestication and global expansion. Nature Genetics 56

54. Yu P, White P, Hochholdinger F, Li CJ. 2014. Phenotypic plasticity of the maize root system in response to heterogeneous nitrogen availability. Planta 240: 667–678

55. Zhan A, Lynch JP. 2015. Reduced frequency of lateral root branching improves N capture from low-N soils in maize. Journal of Experimental Botany 66: 2055–2065

56. Zhan A, Schneider H, Lynch JP. 2015. Reduced Lateral Root Branching Density Improves Drought Tolerance in Maize. Plant Physiology 168: 1603–1615

57. Zhang M, Chen YH, Xing HY, et al. 2023. Positional cloning and characterization reveal the role of a miRNA precursor gene *ZmLRT* in the regulation of lateral root number and drought tolerance in maize. Journal of Integrative Plant Biology 65: 772–790

58. Zhang N, Yu H, Yu H, et al. 2018a. A Core Regulatory Pathway Controlling Rice Tiller Angle Mediated by the *LAZY1*-Dependent Asymmetric Distribution of Auxin. Plant Cell 30: 1461–1475

59. Zhang T, Li RN, Xing JL, Yan L, Wang RC, Zhao YD. 2018b. The YUCCA-Auxin-WOX11 Module Controls Crown Root Development in Rice. Frontiers in Plant Science 9

60. Zhang ZH, Zhang X, Lin ZL, Wang J, Xu ML, Lai JS, Yu JM, Lin ZW. 2018c. The genetic architecture of nodal root number in maize. Plant Journal 93: 1032–1044

61. Zhao Y, Hu YF, Dai MQ, Huang LM, Zhou DX. 2009. The WUSCHEL-Related Homeobox Gene *WOX11* Is Required to Activate Shoot-Borne Crown Root Development in Rice. Plant Cell 21: 736–748

62. Zhou SL, Jiang W, Long F, Cheng SF, Yang WJ, Zhao Y, Zhou DX. 2017. Rice Homeodomain Protein WOX11 Recruits a Histone Acetyltransferase Complex to Establish Programs of Cell Proliferation of Crown Root Meristem. Plant Cell 29: 1088–1104

